# Allergen specific Treg upregulated by lung-stage schistosome infection alleviates allergic airway inflammation via inhibiting IgE secretion

**DOI:** 10.1101/2020.04.14.040998

**Authors:** Zhidan Li, Wei Zhang, Fang Luo, Jian Li, Wenbin Yang, Bingkuan Zhu, Qunfeng Wu, Xiaoling Wang, Chengsong Sun, Yuxiang Xie, Bin Xu, Zhaojun Wang, Feng Qian, Yanmin Wan, Wei Hu

## Abstract

Schistosome infection showed protective effects against allergic airway inflammation (AAI). However, controversial findings exist especially regarding the timing of helminth infection and the underlying mechanisms. Moreover, most previous studies focused on understanding the preventive effect of schistosome infection on asthma (infection before allergen sensitization), while its therapeutic effects (infection after allergen sensitization) were rarely investigated. In this study, we investigated the therapeutic effects of schistosome infection on AAI using a mouse model of OVA induced asthma. To explore how the timing of schistosome infection influences its therapeutic effect, the mice were percutaneously infected with cercaria of *Schistosoma japonicum* at either 1 day before OVA induced asthma attack (infection at lung-stage during AAI) or 14 days before OVA induced asthma attack (infection at post lung-stage during AAI). We found that lung-stage schistosome infection significantly ameliorated OVA-induced AAI, whereas post lung-stage infection showed no therapeutic effect. Mechanistically, the lung-stage schistosome infection significantly upregulated the frequency of Treg, especially OVA specific Treg, in lung tissue, which negatively correlated with the level of OVA specific IgE. Depletion of Treg *in vivo* counteracted the therapeutic effect. Furthermore, transcriptomic analysis of lung tissue showed that lung-stage schistosome infection during AAI shaped the microenvironment to favor Treg induction. In conclusion, our data showed that lung-stage schistosome infection could relieve OVA induced asthma in a mouse model. The therapeutic effect was mediated by the upregulated OVA specific Treg which suppressed IgE production and Th2 cytokine secretion. Our results may facilitate the discovery of a new therapy for AAI.

**Author Summary:** Asthma is an increasingly common disease especially in industrialized countries, which is still lack of an optimal therapy. The protective effect of schistosome infection against allergic asthma has been shown in previous studies, which represents a promising candidate immune intervention approach. However, controversial findings exist especially regarding the timing of helminth infection and the underlying mechanisms. In this study, we demonstrate that lung-stage schistosome infection could upregulate the frequency of allergen specific Treg, which significantly alleviated OVA induced allergic airway inflammation via inhibiting the production of IgE and Th2 cytokines. Our results proved the therapeutic effect of schistosome infection on allergic asthma. Moreover, we highlighted that lung-stage infection is essential for inducing allergen specific regulatory T cells in lung, which is the key mediator of the observed therapeutic effect. These findings shed new light on exploiting helminths or their derivatives to treat asthma and other allergic diseases.

## Introduction

The prevalence of asthma has increased dramatically in the past three decades [1, 2], which represent a great health burden especially in developed countries [3, 4]. Atopic asthma is the most common form of asthma, which is an immunological disorder disease characterized by inflammation of the airways and lungs triggered by allergen with marked Th2 responses, overactive immunoglobulin IgE production, mucus hypersecretion and large amount of eosinophils influx to airways [5].

The exact social and environmental factors that lead to hyper-reactive immune disorder is still not fully understood. A leading theory behind the rapid rising of allergy and asthma rates is the “hygiene hypothesis”, which suggests that the decreasing incidence of infections in western countries is the origin of the increasing incidence of both autoimmune and allergic diseases [6]. The hypothesis was supported by an observation showing that westernized lifestyle linked with significantly higher prevalence of atopic disease [7]. A putative explanation to this phenomenon is that the overall reduction in common Th1-inducing (bacterial, viral and parasitical) infections resulting in a decreased ability to counterbalance Th2-polarized allergic diseases [8-10]. Following this lead, a variety of experimental studies have proved that helminth infection can down-regulate host immunity and immunopathology in allergy and other immune disorders[11-13]. Schistosome was one of the parasites that has been found to have protective effects for autoimmune diseases and allergies like arthritis and asthma [14-16]. These explorations hold great promise to identify a new and better therapy for atopic asthma, which may avoid the adverse effects of current treatments [17-19].

Schistosome is an ancient parasite affecting more than 230 million people in 78 tropical and subtropical countries [20]. During the life stages in the definitive hosts, the trematode invades its mammalian hosts through the skin firstly, migrates from skin to lung, then develops and matures in liver, finally resides mesenteric venules. Although it has been shown by multiple studies that schistosome could abate allergic airway inflammation (AAI), the understanding of underlying mechanisms remains limited. Most previous studies focused on testing the preventive effect (infection before allergen sensitization) of schistosome infection against allergic asthma. And under this setting, controversial results have been reported regarding both the timing of infection (acute versus chronic) [21-23] and the effector component (egg versus worms) [24-26], which reflects the complexities of schistosome life cycle and its immune regulatory components. Moreover, contradictory results were also reported regarding the roles of regulatory T cells in schistosome mediated protection. Some studies showed that Treg was an important effector in schistosome mediated protection against asthma [21, 23, 26-28], while a more recent study showed that the protection was independent of Treg [24].

Unlike previous studies which focused on testing the preventive effect (infection before allergen sensitization) of schistosome infection against allergic asthma, the primary goal of this study was to investigate the therapeutic effect of schistosome infection on asthmatic inflammation (infection after allergen sensitization) and to clarify the underlying mechanism. To this aim, the mice were percutaneously infected with cercaria of *Schistosoma japonicum* at either 1 day before OVA induced asthma attack (infection at lung-stage during AAI) or 14 days before OVA induced asthma attack (infection at post lung-stage during AAI). We found that only lung-stage schistosome infection could upregulate the frequency of allergen specific Treg, which significantly alleviated AAI via inhibiting IgE production and inflammatory cytokine secretion.

## Results

### Lung-stage schistosome infection ameliorated OVA-induced AAI in a murine model

A mouse model of OVA-induced AAI was adopted to test the therapeutic effect of schistosome infection on allergic asthma (Fig 1A & 1B). Compared to the control group, mice in the OVA group showed significant infiltration of inflammatory cells in BALFs (Fig 1C & 1D), which resembled the main clinical feature of AAI [29]. Moreover, after schistosome infection, the results showed lung-stage infection significantly reduced the infiltration of inflammatory cells, especially eosinophils (Fig 1C), while post lung-stage infection did not (Figure 1D). Histopathological examination further confirmed the above findings by showing that lung-stage infection significantly suppressed the OVA-induced eosinophil-rich leukocyte infiltration and mucus hypersecretion (Fig 1E), whereas post lung-stage infection showed no obvious therapeutic effect (Fig 1F).

**Fig. 1.**
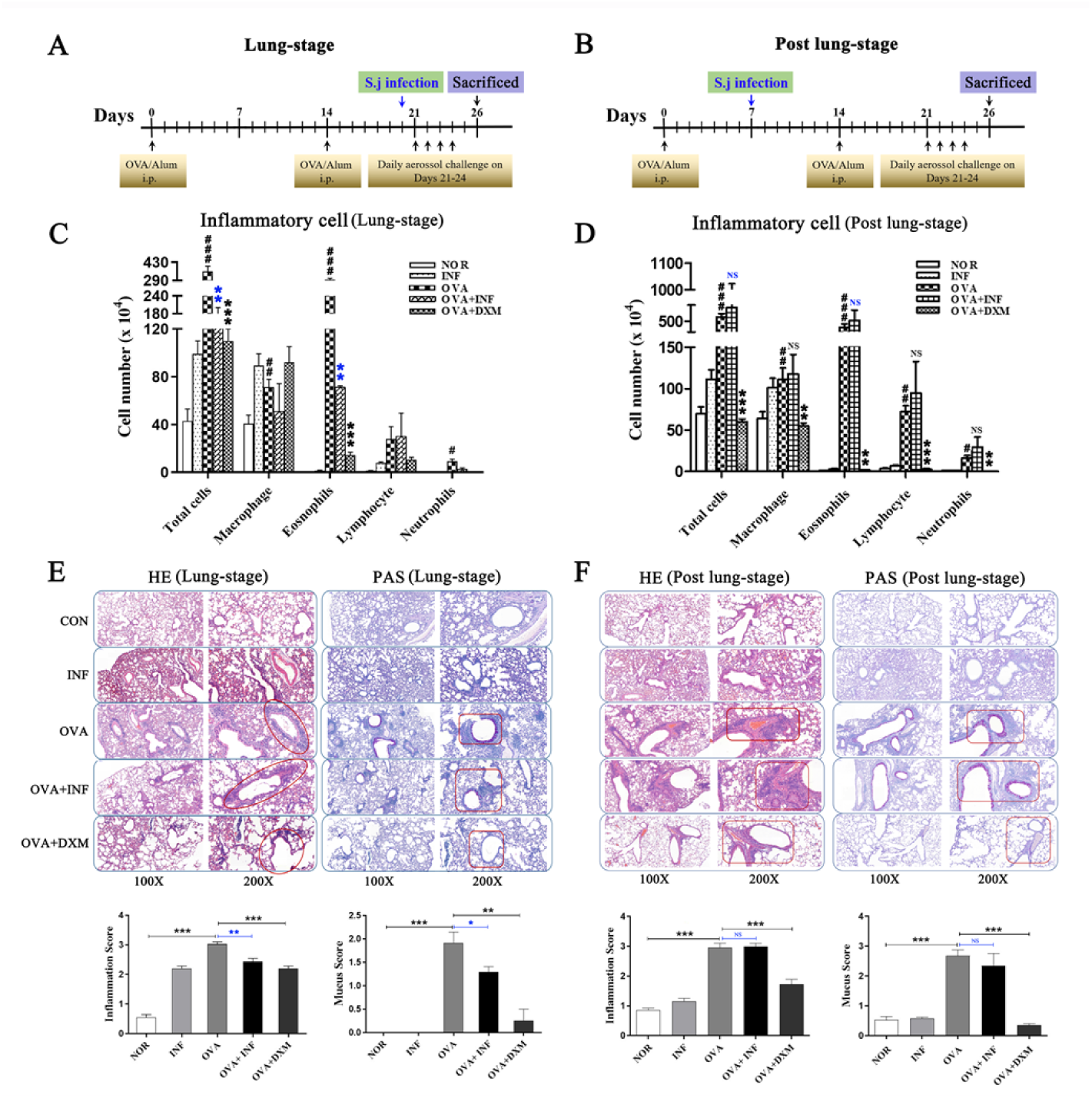
Lung-stage schistosome infection alleviated the attack of OVA-induced AAI, whereas post lung-stage infection did not. Experimental design of OVA induced AAI treated with either lung-stage (**A**) or post lung-stage (**B**) schistosome infection. (**C & D**) Comparisons of inflammatory cell infiltration in BALF of mice after OVA challenge. (**E & F**) Representative images of H&E and PAS staining of lung tissue after OVA challenge. Statistical analysis of inflammation score and mucus secretion score were also shown in (**E**) and (**F**), respectively. NOR, normal mice (without OVA sensitization and challenge); INF, mice without OVA sensitization and challenge but infected with schistosome; OVA, mice with OVA sensitization and challenge but without schistosome infection; OVA + INF, mice sensitized and challenged with OVA and treated with schistosome infection; OVA + DXM, mice sensitized and challenged with OVA and treated with dexamethasone. Data were shown as mean ± SEM, n = 5. *, *P* < 0.05; **, *P* < 0.01; NS, not significant by the one-way analysis of variance (ANOVA) with Tukey test. #; ##; ### indicated *P* < 0.05; < 0.01; < 0.001, respectively, OVA versus NOR (C & D). *, **, *** indicated *P* < 0.05; < 0.01; < 0.001, respectively, OVA+INF or OVA+DXM versus OVA (C & D).

### Lung-stage schistosome infection inhibited IgE production and suppressed Th2 cytokine secretions

IgE is the key factor mediating the pathological immune responses that lead to allergic asthma [30]. To further characterize the therapeutic effects of schistosome infection, we measured the total and OVA specific IgE in serum of mice. The results showed that lung-stage infection significantly downregulated both the total and OVA specific IgE to levels comparable with DXM treated mice (Fig 2A & 2B). In contrast, post lung-stage infection tended to elevate the total and OVA specific IgE levels despite no significant difference was reached (Fig 2C & 2D). Moreover, we also measured a panel of cytokines and chemokines in BALFs and found that lung-stage infection altered the cytokine/chemokine secretion pattern induced by aerosolized OVA challenge (Fig 3A & S1 Fig). More specifically, IL-5 and Eotaxin were reduced to levels similar with DXM treatment (Fig 3B). On the contrary, post lung-stage infection increased IL-4 and IL-5 secretion (Fig 3B).

**Fig. 2.**
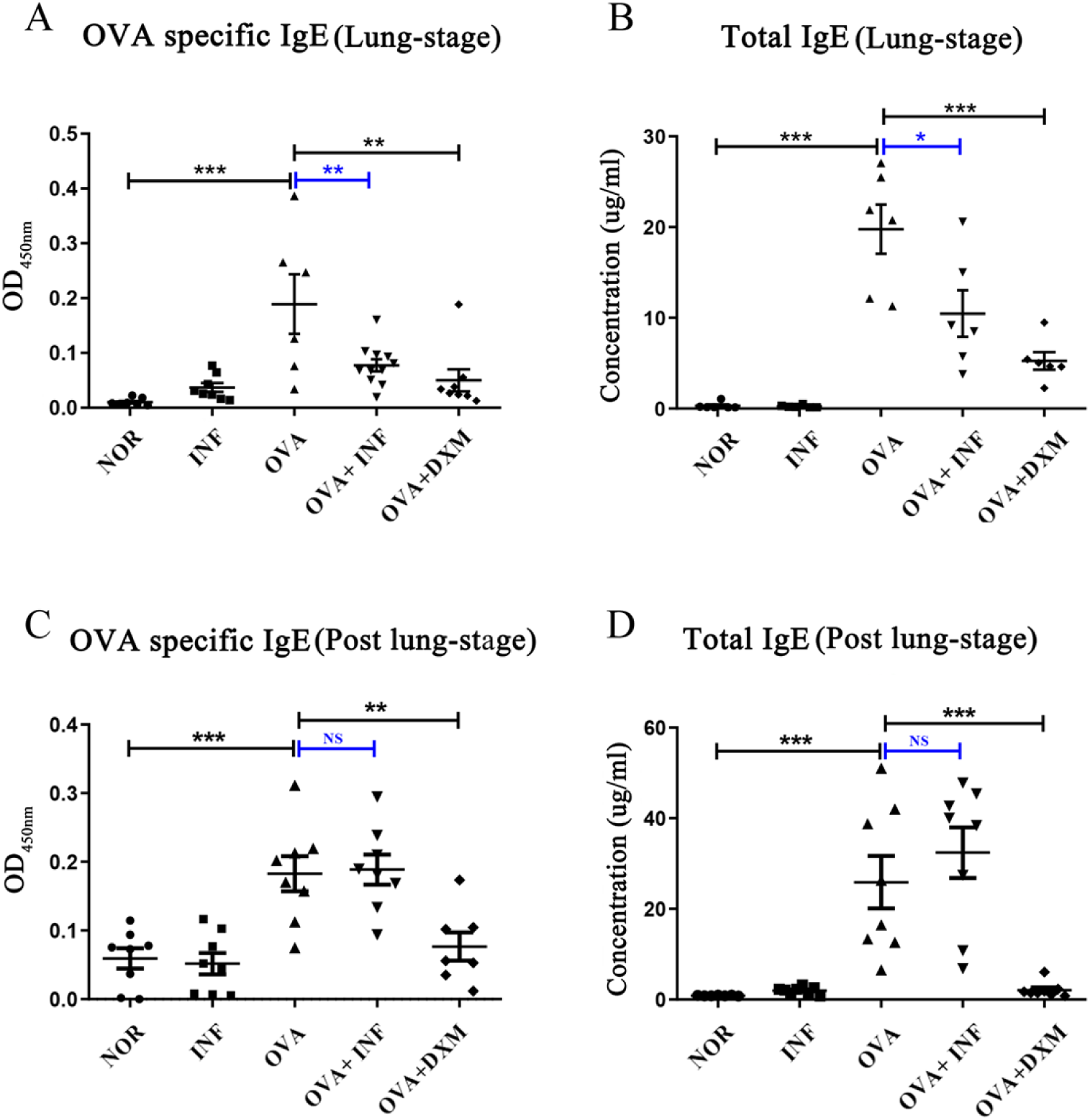
Lung-stage schistosome infection suppressed both the total and OVA specific IgE after OVA challenge, whereas post lung-stage infection did not. (**A & C**) OVA specific IgE in each group were measured by ELISA after treatment with schistosome infection. (**B & D**) The concentration of total IgE in mouse serum were compared among all groups after OVA challenge. Data were shown as Mean ± SEM, n = 5. *, *P* < 0.05; **, *P* < 0.01; ***, *P* < 0.001; NS, not significant.

**Fig. 3.**
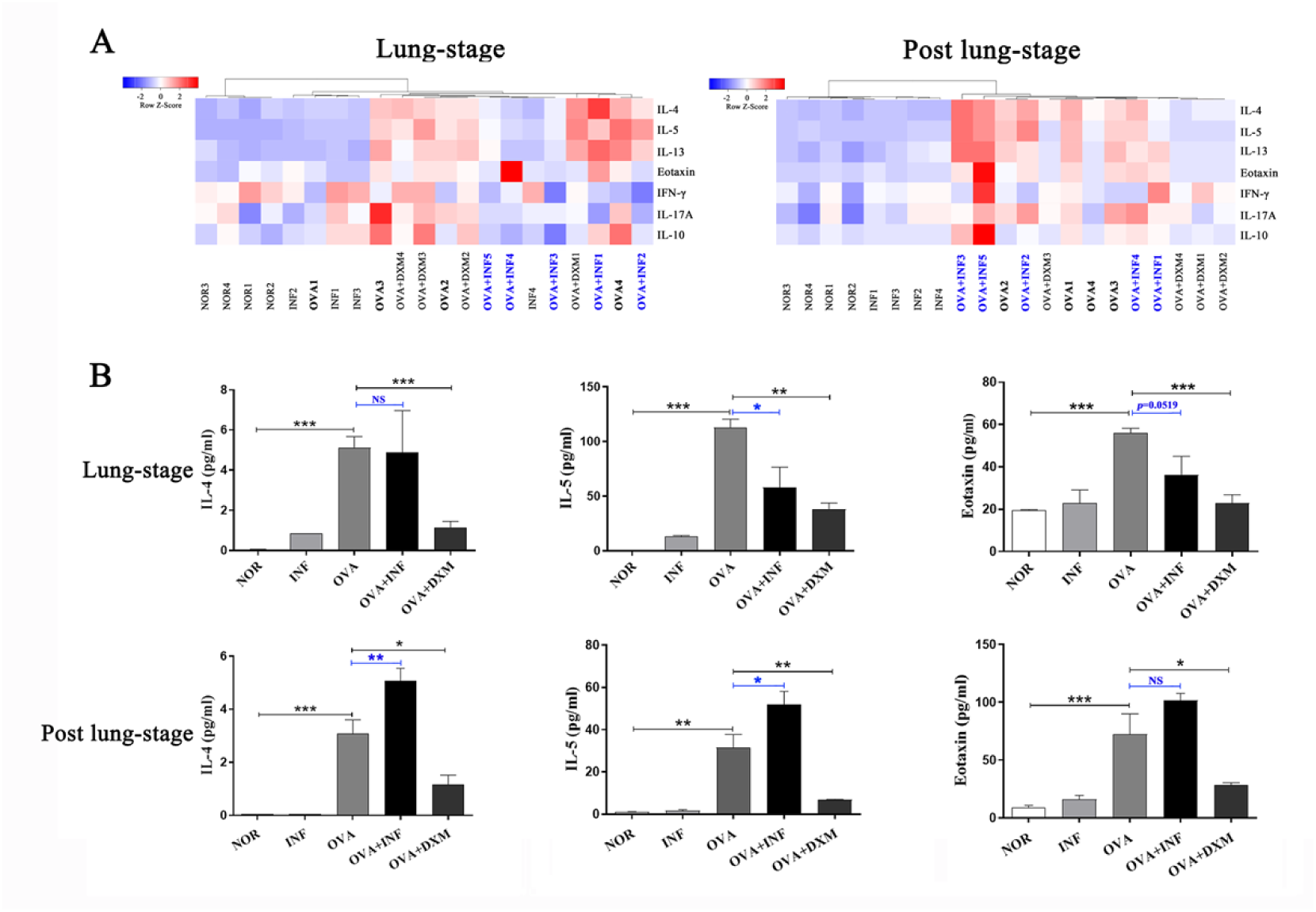
Lung-stage schistosome infection inhibited Th2 cytokine secretion after OVA challenge, while post lung-stage infection did not. **(A)** Heatmaps of multiple cyto-/chemokines of mice treated with lung-stage schistosome infection (left) and post lung-stage schistosome infection (right) after OVA challenge. (**B**) Concentrations of IL-4, IL-5 and Eotaxin in BALFs were compared among all groups. Data were shown as Mean ± SEM, n = 5. *, *P* < 0.05; **, *P* < 0.01; ***, *P* < 0.001; NS, not significant.

### Lung-stage schistosome infection upregulated the frequencies of regulatory T cells (Treg) especially OVA specific Treg in lung

Treg was suggested to be the key factor of *S. mansoni*-mediated protection against allergic airway inflammation [21]. Herein, we first assessed the frequencies of Treg (CD4^+^CD25^+^Foxp3^+^ Treg) in spleen and lung. As shown in Figure 4A, compared to the OVA control, lung-stage infection significantly upregulated the frequency of Treg both in lung and spleen (Fig 4A), whereas post lung-stage infection only slightly improved the proportion of Treg in spleen (Fig 4B). To further illustrated that the influences of schistosome on OVA induced AAI were allergen specific or non-specific immune response, OVA specific naïve CD45.1^+^ CD4^+^ T cells were transferred into CD45.2^+^ recipient mice. The frequencies of total Treg, CD45.1^+^ Treg (OVA specific), and CD45.2^+^ Treg (OVA non-specific) were detected in lung and lung draining lymph nodes (LDLNs). Interestingly, we found that the frequency of OVA specific Treg (CD45.1^+^ Treg) in lung increased by more than 3 folds after schistosome infection (*P* < 0.001), while that in LDLNs didn’t show any significant changes (Fig 5B & 5C). However, the frequency of endogenous Treg (CD45.2^+^ Treg) in lung was not significantly improved, while that in LDLNs showed a slight increased (Fig 5B & 5C). The proportion of total Treg was increased in lung and LDLNs (Fig 5B & 5C).

**Fig. 4.**
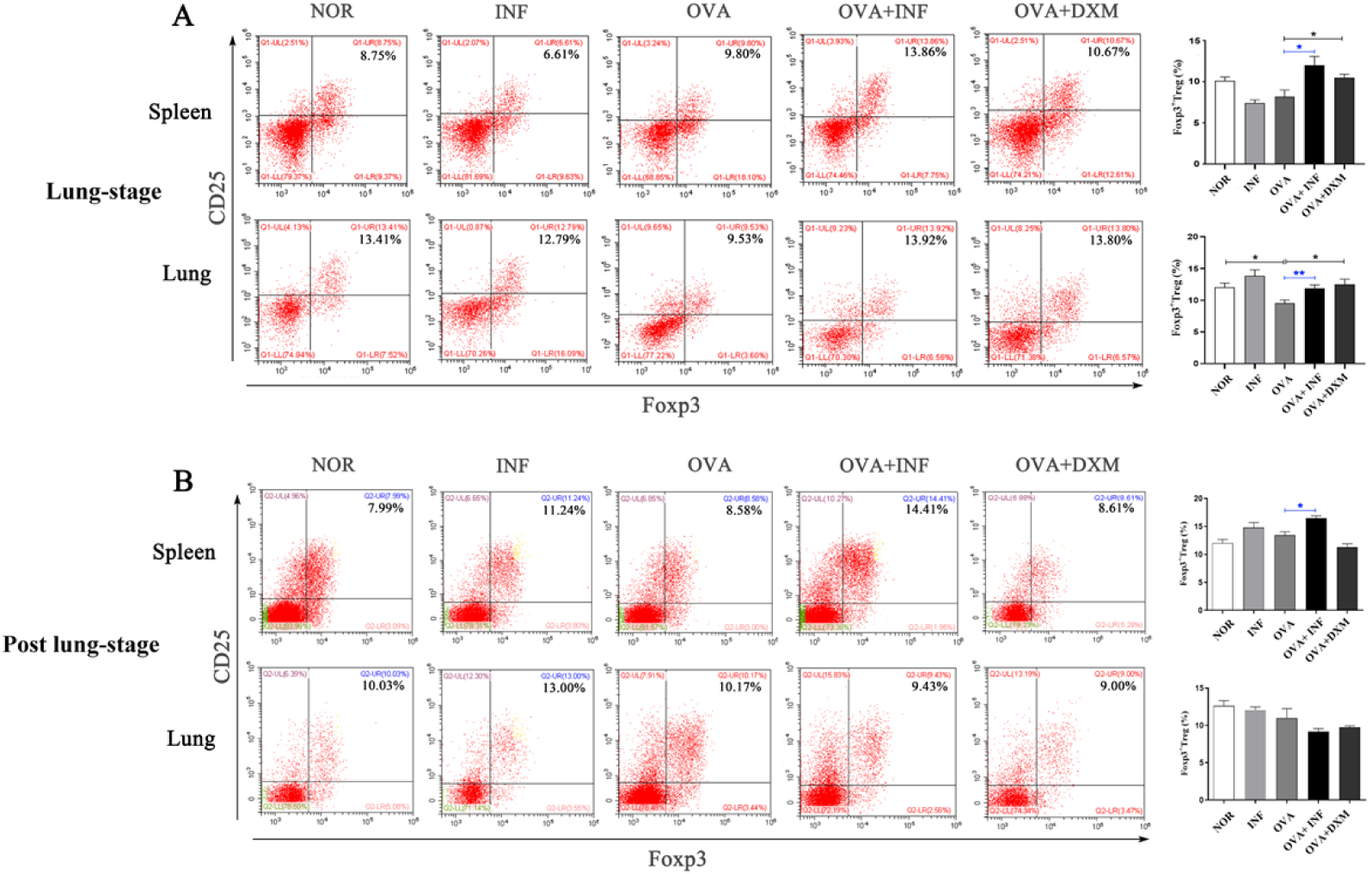
Lung-stage schistosome infection upregulated Treg frequency in lung and spleen after OVA challenge. (**A & B**) Comparisons of Treg frequencies (CD4^+^CD25^+^Foxp3^+^ Treg) in lungs and spleens among all groups. Representative data of flow cytometry analysis for each group were shown together with statistical comparisons. Data were presented as Mean ± SEM, n = 5. *, *P* < 0.05; **, *P* < 0.01; ***, *P* < 0.001.

**Fig. 5.**
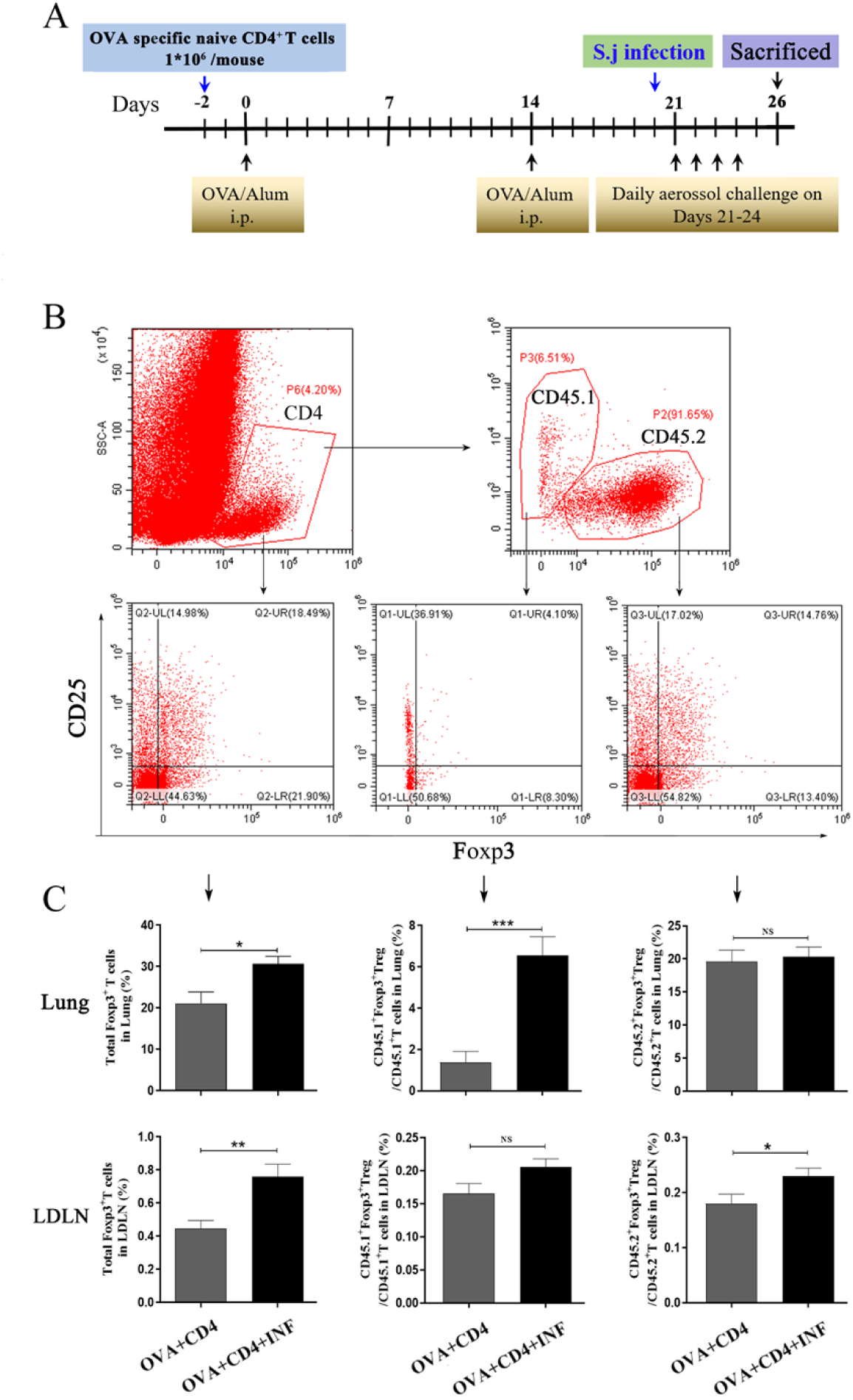
Lung-stage schistosome infection upregulated OVA specific Treg after OVA challenge. **(A)** Design of experiment for testing the therapeutic effect of lung-stage schistosome infection on OVA induced AAI after adoptive transfer of OVA specific naïve CD4^+^ T cell. (**B & C**) Gate strategy and statistical comparisons of flow cytometry analysis for total Treg, CD45.1^+^ Treg (OVA specific) and CD45.2^+^ Treg in lung and lung draining lymph nodes (LDLN). Data were shown as Mean ± SEM, n = 8. *, *P* < 0.05; **, *P* < 0.01; ***, *P* < 0.001; NS, not significant.

Besides, we also found that the ratio of OVA specific CD4^+^ IL-4^+^ T versus CD4^+^ IFN-γ^+^ T cells significantly decreased after lung-stage schistosome infection (S2 Fig), suggesting that specific CD4^+^ T cell responses from Th2 toward Th1 shifted responses.

### The therapeutic effect of lung-stage schistosome infection was Treg dependent

Significant negative correlations between the frequency of Treg and OVA specific IgE or IgG (Fig 6) were observed, indicating that the therapeutic effect of schistosome infection on AAI might be mediated by Treg. To elucidate the role of Treg, we performed *in vivo* depletion using anti-mouse CD25 antibody (Fig 7A). Our data showed that Treg depletion (OVA+INF+αCD25 group) aggravated OVA induced AAI compared to isotype control group. Inflammatory cell infiltration, mucus secretion (shown by PAS staining) and OVA specific IgE production significantly increased after Treg depletion (Fig 7).

**Fig. 6.**
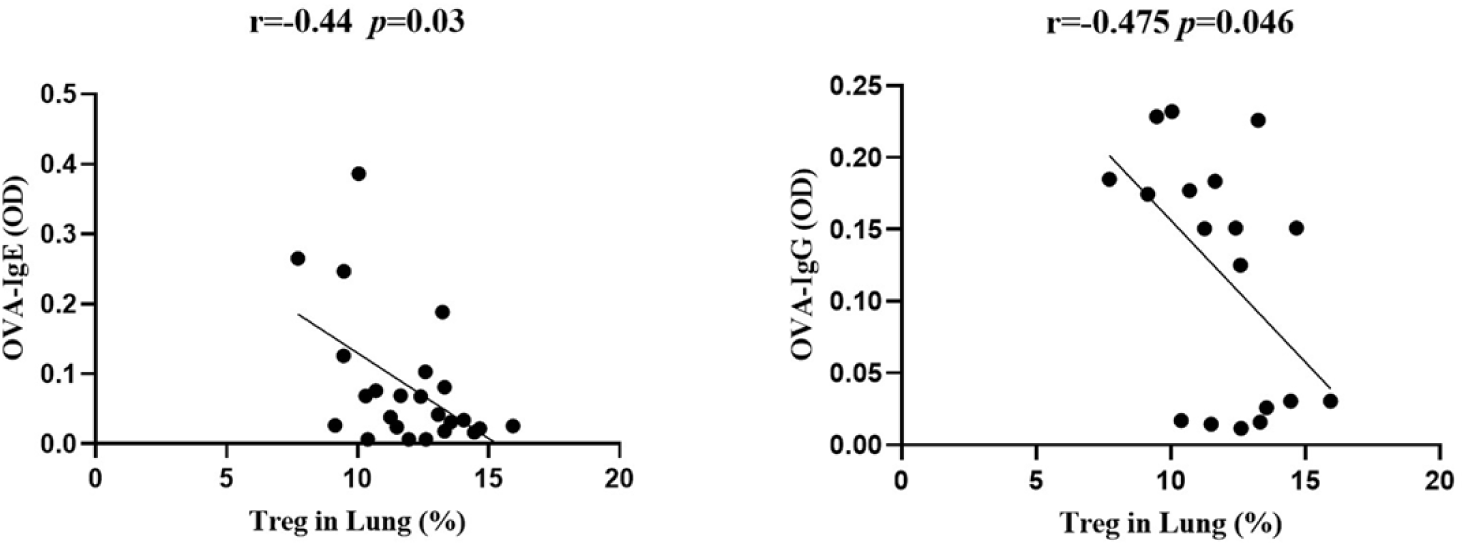
The frequency of Treg in lung negatively correlated with OVA specific IgE and IgG. Correlation analysis between Treg frequency in lung and the OD values of OVA specific IgE (left) and IgG (right) in serum.

**Fig. 7.**
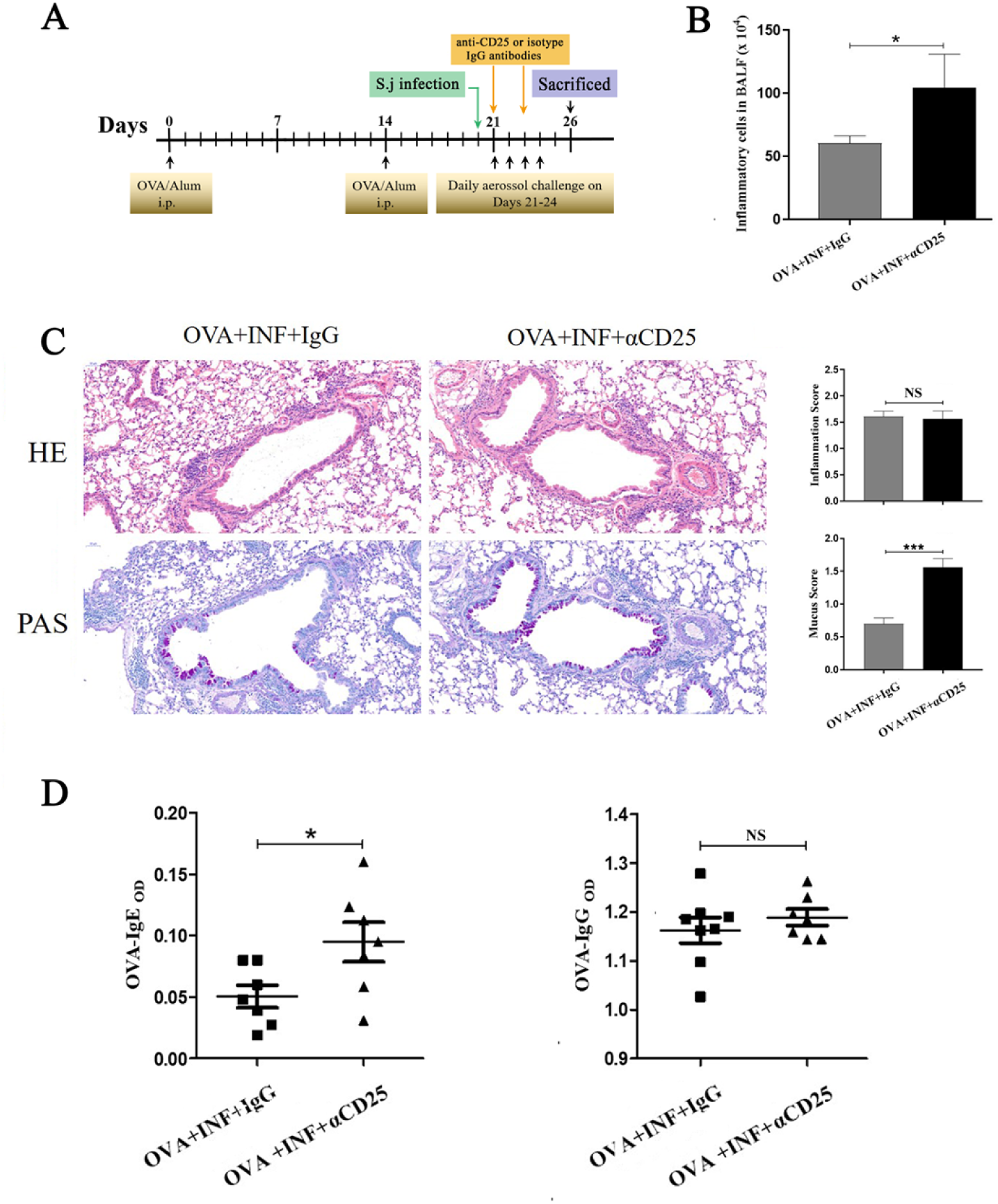
*In vivo* depletion of Treg counteracted the therapeutic effect of lung-stage schistosome infection on OVA-induced AAI. (**A**) Design of experiment for testing the role of Treg in the therapeutic effect mediated by lung-stage schistosome infection. (**B**) Comparisons of inflammatory cell counts in BALF between lung-stage schistosome infected mice treated with either anti-CD25 antibody or isotype control IgG. (**C**) Lung histopathology analysis of lung-stage schistosome infected mice treated with either anti-CD25 antibody or isotype control IgG. Upper, H&E staining; lower, PAS staining. (**D**) Comparisons of OVA specific IgE and IgG between Treg depleted and control mice. Data were shown as Mean ± SEM, n = 8. *, *P* < 0.05; ***, *P* < 0.001.

### Lung-stage schistosome infection moulded the microenvironment to facilitate the generation of Treg

To find out factors that contributed to the induction of Treg upon lung-stage schistosome infection, we performed the transcriptomic profiles of the lung tissues from the schistosome infected and non-infected mice post OVA challenge. The results showed that 203 genes were upregulated and 279 genes were downregulated in the lung-stage schistosome infection group (Fig 8A & Data file S1). GO analysis of DEGs showed that the top 3 terms of significantly enriched (*P* < 0.05) is mainly distributed in the T cell activation, the leukocyte proliferation and the regulation of leukocyte proliferation (Fig 8B) pathways. And panther analysis showed that 84 DEGs are relate to immune system process (S3 Fig) and 70 of them were downregulated (Data file S1). Further analysis showed that 3 genes (CD46, Epor, and Klra17) reported to promote Treg response were upregulated [31-33] and 8 genes (Clec7a, CCR6, Spi-B, ABCG1, ADA, Ctsk, Ctss, and Ptgir) reported to inhibit Treg response were downregulated [34-41] (Fig 8C & Table 1) in schistosome infected mice. We postulated that lung-stage schistosome infection generated a microenvironment facilitating Treg development in lung (Fig 8C).

**Table 1.**
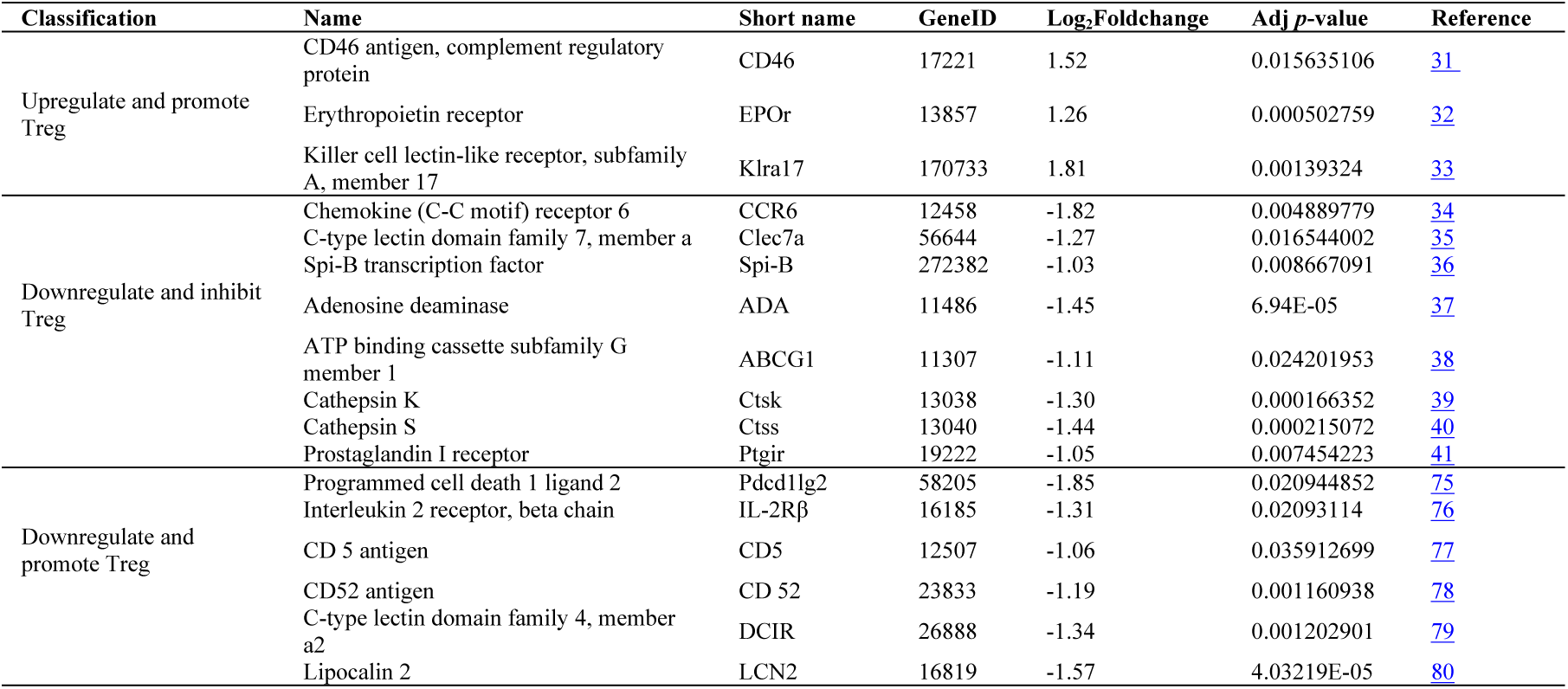
DEGs reported to promote or inhibit Treg response.

**Fig. 8.**
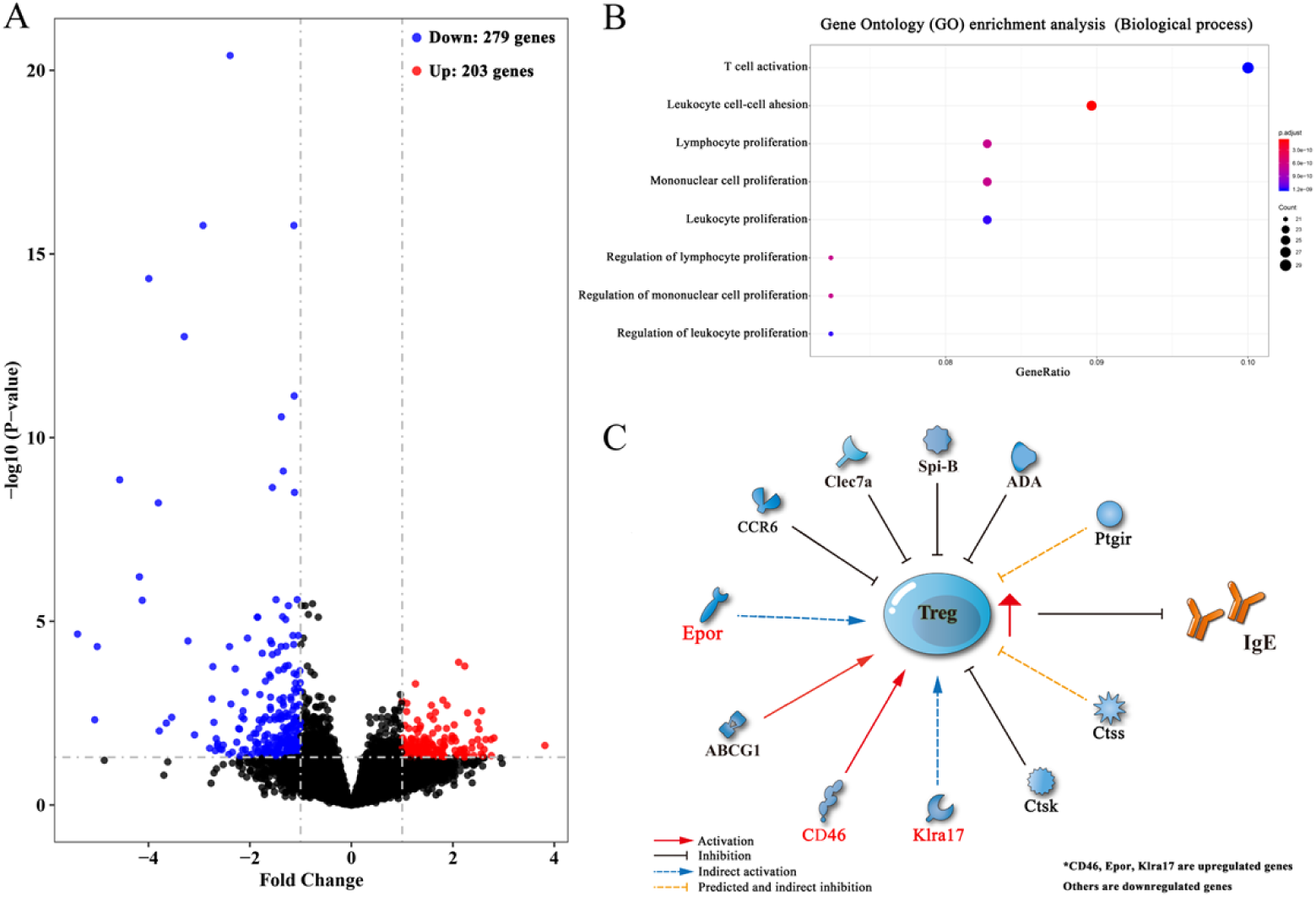
Transcriptomic analysis of differentially expressed genes (DEGs) between lung tissues of OVA-induced asthmatic mice treated with and without lung-stage schistosome infection. (**A**) Volcano plot of detected gene transcription profile in lung tissues of OVA-induced asthmatic mice treated with lung-stage schistosome infection compared with no-treatment control mice after OVA challenge. (**B**) The top 8 functional enrichment pathways of Gene ontology (GO) analysis for biological process in DEGs (*P* < 0.05). (**C**) Predicted gene network that might promote the generation of Treg in DEGs.

In addition, we found that 8 genes (DOCK2, IRF4, Rac2, Lgals3, H2-Oa, Pdcd1lg2, Sash3, and Mzb1) related to B cell function or differentiation [42-49] were also downregulated after schistosome infection (Table 2), which might potentially contribute to the inhibition of IgE response. Genes related to lung development (FOXF1, ANO9, TRIM6, MMP27, Epor, Gata1, and Serpina) [50-52] and cell integrity (Villin and CRB1)[53, 54] were found to be upregulated too, which indirectly supported the observed therapeutic effect of schistosome infection (Table 2).

**Table 2.**
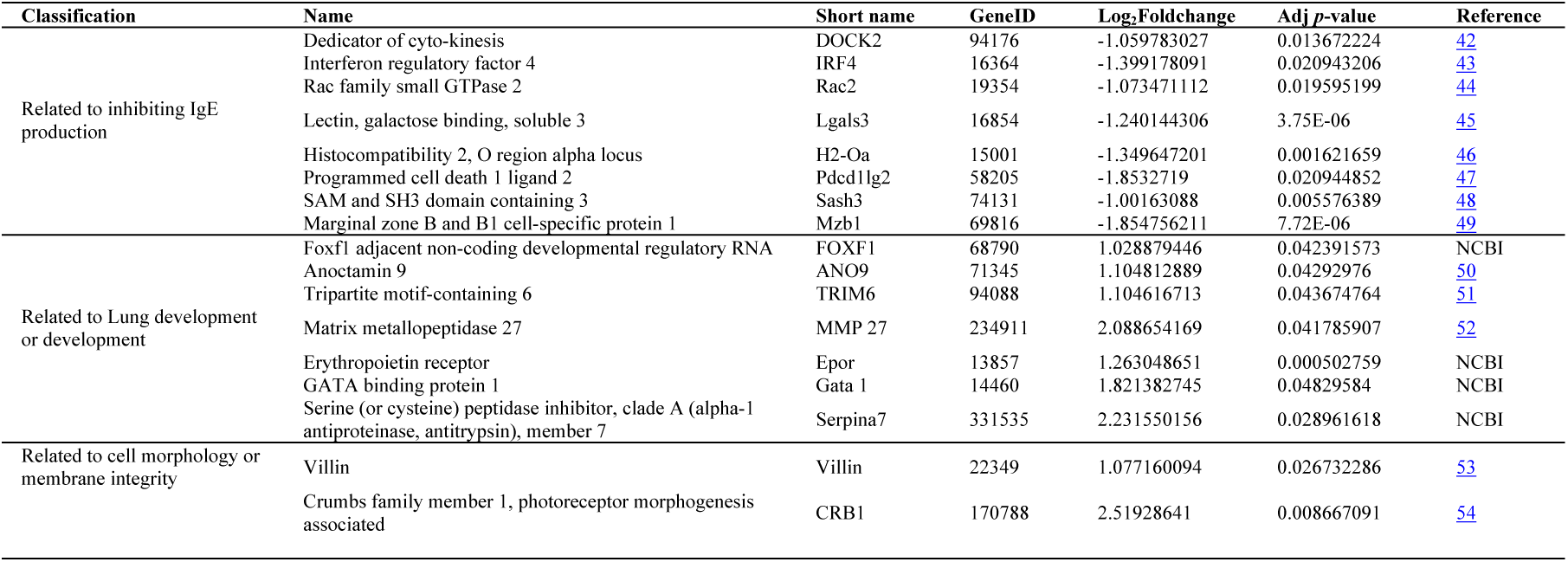
DEGs reported to facilitate B cell or plasma cell, lung development and cellular morphology.

## Discussion

The eradication of helminths (and other pathogens) is suggested to have resulted in over-activated immune response, which might be the cause of the increasing prevalence of allergic and autoimmune disorders especially in developed and urbanized countries [55-57]. The therapeutic effect of parasitic infection against allergies and autoimmune disease have been extensively explored especially after the hygiene hypothesis was introduced into this field [58]. Among which, the immunoregulation of schistosome is best illustrated [14, 21, 59].

In this study, to investigate how the timing of schistosome infection influence the development of allergic asthma, we compared the therapeutic effect of two phase of schistosome infection: lung-stage and post lung-stage. We found that lung-stage schistosome infection significantly relieved OVA-induced allergic airway inflammation, but post lung-stage infection showed no therapeutic effect. Within lung-stage infection (3-7 days post infection), schistosomula transformed from cercaria was completely located in lung tissue of the host [60], which might modulate the local immune response to abate OVA induced AAI. We postulated that this might be the reason that made the therapeutic effect of lung-stage infection superior to post lung-stage infection. And indeed, we found that lung-stage infection significantly upregulated Treg response in lung.

Multiple factors such as worm species, timing, intensity and chronicity of infection, as well as host genetics have been investigated to illustrate the mechanisms of helminth mediated the regulation of host immunity [61]. Nonetheless, the relationship between helminths and asthma still remains. Mechanistic studies reported contradictory results, for example, one study showed that *S. mansoni*-mediated suppression of allergic airway inflammation was patency dependent and mediated by infection-induced Treg [21], while another study showed that protection mediated by *S. mansoni* egg was independent of either Treg or Breg [24]. In current study, we found that lung-stage schistosome infection occurred during OVA induced asthma attack could upregulate the frequency of Treg and suppressed OVA specific IL-4 response. Upregulation of Treg by schistosome infection has been reported by few previous studies [21, 62], however, to our knowledge, this is the first proof showing that the lung-stage schistosome infection can upregulate allergen OVA specific Treg.

To elucidate the role of Treg in schistosome infection mediated alleviation of AAI, we first analyzed the relationship between Treg and OVA specific IgE and found that the frequency of Treg in lung negatively correlated with OVA specific IgE. Furthermore, by *in vivo* depletion of Treg, we found that the decrease of IgE secretion was Treg dependent. IgE acts as the major mediator resulting in the allergic airway inflammation [63]. Our result proved that the therapeutic effect of schistosome infection on AAI was mediated by a Treg dependent inhibition of IgE, which was consistent with a previous report showing that the preventive effect of chronic *S. mansoni* infection against later AAI was also Treg dependent [21].

Mechanisms underlying the induction of Treg or Breg by egg related antigens have been reported [64, 65]. However, we did not find out the exact active molecules of schistosome that led to the upregulation of Treg in this study. Nonetheless, we think that it is very likely the observed therapeutic effect was a collective result of multiple components of schistosome, as previous studies showed multiple enzymes released by schistosomula could regulate host immunity [66, 67]. We plan to acutely define these components in future.

Instead of identifying effector antigens, in this study, we tried to understand how the lung-stage schistosome infection influence local immune responses in lung. To do so, we performed transcriptomic comparison between lung tissues of schistosome infected and non-infected mice. The results showed, after lung-stage schistosome infection, most genes related to immune response were downregulated (70/84), implying the general immune state in lung tended to be downregulated by schistosome infection. Among these genes, we found that 3 genes (CD46, Epor, and Klra17) reported to promote Treg response were upregulated and 8 genes (Clec7a, CCR6, Spi-B, ABCG1, ADA, Ctsk, Ctss, and Ptgir) reported to inhibit Treg response were downregulated in schistosome infected mice, suggesting that schistosome infection generated a milieu facilitating Treg induction in lung. In the meantime, we also observed some molecules reported to facilitate the function of B or plasma cells were downregulated, which was consistent with our finding that IgE response was suppressed.

Collectively, our study showed that lung-stage schistosome infection established a regulatory environment in lungs, which can help to relieve OVA induced AAI in mouse model. Although the exact mechanism about Treg upregulation remains elusive, our data clearly showed that lung-stage schistosome infection can improve the frequency of allergen specific Treg and the latter can directly suppress IgE production. The encouraging results highlight the value of lung-stage schistosome infection as a potential therapy for allergic asthma. And identifying the effector molecules is especially of interesting, as it will make this therapy more practical.

## Methods

### OVA-induced allergic airway inflammation and schistosome infection

Female BALB/c mice (6- to 8-week-old) were randomly divided into six groups in this experiment, which are OVA-induced AAI (OVA) group, OVA-induced AAI with lung-stage schistosome infection (OVA+INF, lung-stage) group, OVA-induced AAI with post-lung stage infection (OVA+INF, post lung-stage) group, OVA-induced AAI with dexamethasone (DXM) treatment (OVA+DXM) group, as well as infection (INF) group and normal (NOR) group. The mice were sensitized by injecting 10 µg of alum-adjuvanted ovalbumin (OVA; Cat# 77120 and 77161, Thermo Fisher, US) intraperitoneally on day 0 and day 14. Subsequently, to induce AAI, the mice were challenged with aerosolized OVA (1% in PBS) for 30 minutes in the chamber of a Medical Compressor Nebulizer (DEDAKJ, Germany) on days 21–24 (Fig 1A and 1B). The mice of the normal control and schistosome infection control groups were challenged with phosphate buffer solution (PBS). To test the therapeutic effect of infection on OVA induced AAI, mice were infected with 15 cercaria of *S. japonicum* at either 1 day before OVA induced asthma attack (infection at lung-stage during AAI) or 14 days before OVA induced asthma attack (infection at post lung-stage during AAI).

### Bronchoalveolar lavage collection and cell counting

Mice were euthanized 48 h after the last aerosolized OVA challenge (day 26), and bronchoalveolar lavage fluids (BALFs) were collected as previously reported method [68]. Briefly, after euthanasia, tracheotomy was carried out and an arteriovenous indwelling needle (20G; BRAUN, Germany) was inserted into the trachea. Lavages were collected by washing the lung twice with 0.3 ml PBS. Cells in BALFs were harvested after centrifugation and the supernatants were stored at −80°C for cytokine detection. Cell pellet was fixed with paraformaldehyde (4%) and stained with a Haematoxilin-Eosin (H&E). A total of 1000 cells from multiple fields were examined for each slide. Counts of total cells, eosinophils, macrophage, neutrophils, and lymphocytes were performed on blinded samples, as described previously [69].

### Lung histopathology

Lung tissues were fixed in 4% phosphate buffered formaldehyde overnight, then embedded in paraffin and cut for haematoxylin-eosin (H&E) and periodic acid-Schiff (PAS). Images of the stained sections were captured with a NIKON DS-U3 microscope (NIKON, Japan). Lung inflammation and the intensity of goblet cell metaplasia was assessed and scored 0-4 by two blinded, independent investigators, as described previously [70].

### Determination of total and OVA-specific IgE in serum

The level of total and OVA specific IgE in serum were measured using enzyme linked immunosorbent assay (ELISA). Briefly, Maxisorp 96-well microtiters plates (Thermo Fisher Scientific, USA) were coated with rat monoclonal anti-mouse IgE antibody for total IgE detection (1: 1000; Cat# ab99571, Abcam, UK) or 10 μg/ml ovalbumin for OVA specific IgE (Cat# A5503, Sigma, US) 100 μl/well, respectively, in carbonate-bicarbonate buffer, pH 9.6, for 12–16 hours at 4°C. Then the plates were blocked for at least 2 hours at 37°C with 100 μl/well of PBS plus BSA (1%). After wash, 100 μl serum diluted with PBST (1: 40 for total IgE; 1: 5 for OVA specific IgE) were added to each well and incubated at 37°C for 2 hours. Next, HRP labeled goat anti-mouse IgE antibody were diluted with PBST (1: 2000; Cat# ab99574, Abcam, UK) and added to each well at 100 μl/well. After 2 hours incubation at 37°C, the plates were washed with PBST for 5 times. Finally, color was developed by addition of 100 μl/well of TMB (Cat# PA107, TIANGEN, China) and after incubating at room temperature for maximal 30 minutes, the reaction was stopped with 5% sulfuric acid (50 μl/well). Optical density (OD) values were determined at 450 nm using the multi-mode microplate readers (BioTek, USA). The concentration of total IgE was then calculated according to the standard curve.

### Cytokine detection in BALFs

Levels of IL-4, IL-5, IL-13, IL-10, Eotaxin and IFN-γ in BALFs were measured using a custom-made Bio-Plex Pro Reagent Kit V (6-plex customization) (Cat# MHSTCMAG-70K, Wayen Biotechnologies, China) according to the manufacturer’s instructions. The fluorescence labeled beads was detected using a corrected Bio-Plex MAGPIX system (Bio-Rad, Luminex Corporation, Austin, TX, USA) and the cytokine concentrations were calculated using Bio-plex manager 6.1 (Bio-Rad).

### Lymphocytes isolation from lung tissues

After collection, lung tissues were washed 3–4 times with Roswell Park Memorial Institute (RPMI) medium, minced to tiny pieces, and then digested in 0.1% type IV collagenase (Cat# C8160, Solarbio, China) solution at 37°C for 30 min. Digested lung tissues was filtered through a 70 μm cell strainer and erythrocytes were lysed with a Red Blood Cell Lysis Buffer (Cat# R1010, Solarbio, China).

### Flow cytometry assay

Single cells suspension were stained with a panel of surface mAbs in FACS buffer (PBS containing 2 mM EDTA and 0.5% bovine serum albumin) for 30 min on ice, including FITC-conjugated anti-CD4 (Clone# 88-8111-40, eBioscience, USA), APC-conjugated anti-CD25 mAb (Clone# 88-8111-40, eBioscience, USA), SuperBright645-conjugated anti-CD45.1 (Clone# 64-0453-82, eBioscience, USA) and Pe-cyanine7-conjugated anti-CD45.2 (Clone# 25-0453-82, eBioscience, USA). Subsequently, cells were fixed with fix/perm buffer (Clone# 88-8111-40, eBioscience, USA) on ice for 20 min, and then stained with mAbs targeting intracellular markers in a Perm/wash buffer for 30 min on ice. For the detection of Treg, PE labeled anti-Foxp3 mAb (Clone# 88-8111-40, eBioscience, USA) was used. And for detecting OVA specific IL-4 and IFN-γ secretion, isolated lymphocytes were initially stimulated for 16 h with 5 ug/ml OVA peptide (323-339) (China peptides, China) and then stained with mAbs Perp-cy5.5 conjugated anti-CD3 (Clone# 145-2C11, eBioscience, USA) and FITC conjugated anti-CD4 (Clone# 88-8111-40, eBioscience, USA) for 30 min on ice. Subsequently, cells were fixed with fix/perm buffer (Clone# 88-8111-40, eBioscience, USA) on ice for 20 min. Then PE conjugated anti-IL-4 (Clone# 12-7041-81, eBioscience, USA) or APC conjugated anti-IFN-γ (Clone# 17-7311-81, eBioscience, USA) for 30 min on ice. Finally, after two washes, all cells were resuspended in PBS containing 1% paraformaldehyde and subject to flow cytometry analysis (Cytometer LX, Beckman).

### Adoptive Transfer of naïve CD4^+^ T cells

Naïve CD4^+^ T cells of CD45.1^+^ OT II mice were purified using EasySep Mouse Naïve CD4^+^ T Cell Isolation Kit (Cat# 19765, StemCell, USA) according to the manufacturer’s protocol. The purity of isolated cells was checked by flow cytometry and was confirmed to be > 85%. Freshly purified naïve CD4^+^ T cells were suspended in PBS and injected intravenously into CD45.2^+^ congenic C57BL/6 recipient mice, 1 × 10^6^ cells/mouse. The induction of AAI and schistosome infection were performed as described above.

### *In vivo* depletion of Treg

Anti-CD25 antibody clone PC61 has been widely used to deplete Tregs for characterizing Treg function *in vivo* [71]. 100 μg/mouse anti-CD25 antibody (Cat# 16-0251-85, Clone# PC61.5, eBioscience, USA) or isotype IgG (Cat# 16-4301-85, Clone# eBRG1, eBioscience, USA) were dissolved with 150 μl sterile PBS and injected intravenously into the mice 21 days post OVA sensitization. A second shot of 50 μg /mouse antibodies was given on day 23 post OVA sensitization (Fig 7A). After depletion, the mice were randomly divided into two groups: OVA+INF+αCD25 and OVA+INF+IgG. OVA sensitization, aerosol challenge and schistosome infection were performed as described above.

### RNA sequencing

Total RNA was extracted from lung tissues by using Trizol reagent (Cat# 15596026, Invitrogen). RNA purity was checked using the Nano Photometer spectrophotometer (IMPLEN, CA, USA). RNA integrity was assessed using the RNA Nano 6000 Assay Kit of the Bioanalyzer 2100 system (Agilent Technologies, CA, USA). 1 μg total RNA from each sample was used to construct the sequencing library using Poly(A) mRNA Capture Module (Cat# RK20340, Abclonal, USA) and Fast RNA-seq Lib Prep Module for Illumina (Cat# RK20304, Abclonal, USA). Index codes were added to attribute sequences of each sample. Then the libraries were sequenced on Illumina Novaseq platform (2 × 150 bp). Total 7 samples, 3 from OVA group and 4 from OVA+INF group, were sequenced in one lane, producing more than 30 million reads per library.

### Differential expression genes (DEGs) analysis and functional enrichment analysis

Sequencing quality was evaluated by FastQC software (http://www.bioinformatics.babraham.ac.uk/projects/fastqc/). Poor quality reads and adaptors were trimmed by Trimmomatic software (Released Version 0.22, www.usadellab.org/cms/index/php?page = trimmomatic), and only reads longer than 50 bp were used for further analysis. The high-quality reads were mapped to mouse genome (mouse BALB/cJ) downloaded in Ensembl database. The HTseq [72] were used to quantify gene expression and R DEseq2 package [73] were employed for differential expression analysis. Only genes with FDR adjusted *P*-value < 0.05 and absolute value of fold change > 2 were considered as DEGs. Functional enrichment of GO terms and KEGG analyses of DEGs were conducted by R cluster Profiler package [74] with FDR correction. Significantly enriched GO terms and KEGG pathways were identified with corrected *P* value < 0.05. DEGs related pathways enrichment terms were performed with the Panther Classification System (http://pantherdb.org/).

RNA sequencing data are deposited in the SRA database, SRA accession number: PRJNA609083.

### Ethics Statement

All experiments and methods were performed in accordance with relevant guidelines and regulations. Mice experiments were carried out at National Institute of Parasitic Disease, Chinese Center for Disease Control and Prevention (NIPD, China CDC) in Shanghai, China. All animal experiment protocols used in this study were approved by the Laboratory Animal Welfare & Ethic Committee (LAWEC) of National Institute of Parasitic Diseases (Permit Number: IPD-2016-7).

### Statistical analysis

All statistical analyses were performed using GraphPad Prism 8.0 (GraphPad Software, Inc., San Diego, CA, USA). The data of quantitative variables were presented as mean ± standard error of mean (SEM). *P* < 0.05 was considered statistically significant.

## Supporting information

**S1 Fig.** Comparisons of concentrations of IL-13, IL-10, IL-17A and IFN-γ in BALF. FI indicated fluorescence intensity. Data were shown as Mean ± SEM. *, P < 0.05; **, P < 0.01; ***, P < 0.001.

(TIF)

**S2 Fig.** The influences of Lung-stage schistosome infection on OVA specific IFN-γ and IL-4 response after OVA challenge.

(A) Gating strategy of flow cytometry. (B) Frequencies of OVA specific CD3+CD4+IL-4+ T cells, CD3+CD4+IFN-γ+ T cells and their ratios in lung and LDLN. Data were shown as Mean ± SEM, n = 8. *, P < 0.05; **, P < 0.01 and NS, not significant.

(TIF)

**S3 Fig.** Panther pathway analysis of DEGs between lung-stage schistosome infected mice and no-treatment control mice post OVA challenge.

(TIF)

## Acknowledgements

We thank the staff of snail house at National Institute of Parasitic Disease, Chinese Center for Disease Control and Prevention (NIPD, China CDC) for supporting the cercaria of *Schistosoma japonicum*. We acknowledge the animals that contributed to this study.

## Author contribution

**Conceptualization:** Zhidan Li, Yanmin Wan and Wei Hu.

**Data curation:** Zhidan Li.

**Formal analysis:** Zhidan Li and Fang Luo

**Funding acquisition:** Wei Hu.

**Investigation:** Zhidan Li, Wei Zhang, Fang Luo, Jian Li, Wenbin Yang, Bingkuan Zhu, Qunfeng Wu, Xiaoling Wang, Chengsong Sun, Yuxiang Xie, Bin Xu, Zhaojun Wang, Feng Qian, Yanmin Wan and Wei Hu.

**Methodology:** Zhidan Li, Wei Zhang, Fang Luo, Jian Li, Wenbin Yang, Bingkuan Zhu, Qunfeng Wu, Xiaoling Wang, Chengsong Sun, Yuxiang Xie.

**Project administration:** Yanmin Wan and Wei Hu.

**Resources:** Bin Xu, Zhaojun Wang, Feng Qian, Yanmin Wan and Wei Hu.

**Supervision:** Yanmin Wan and Wei Hu. **Visualization:** Zhidan Li, Yanmin Wan and Wei Hu. **Writing – original draft:** Zhidan Li.

**Writing – review & editing:** Yanmin Wan and Wei Hu.

**Data file S1**. DEGs between OVA and OVA+INF groups and their classifications. (See supplementary materials)

## References

1. Ali FR. Does this patient have atopic asthma? Clin Med (Lond). 2011;11(4):376–80. Epub 2011/08/23. https://doi.org/10.7861/clinmedicine.11-4-376. PubMed PMID: 21853839; PubMed Central PMCID: PMCPMC5873752.

2. Eder W, Ege MJ, von Mutius E. The asthma epidemic. N Engl J Med. 2006;355(21):2226–35. Epub 2006/11/25. https://doi.org/10.1056/NEJMra054308. PubMed PMID: 17124020.

3. Thomsen SF. Epidemiology and natural history of atopic diseases. Eur Clin Respir J. 2015;2. Epub 2015/11/12. https://doi.org/10.3402/ecrj.v2.24642. PubMed PMID: 26557262; PubMed Central PMCID: PMCPMC4629767.

4. Barnes PJ. The size of the problem of managing asthma. Respir Med. 2004;98 Suppl B:S4–8. Epub 2004/10/16. https://doi.org/10.1016/j.rmed.2004.07.009. PubMed PMID: 15481282.

5. Lambrecht BN, Hammad H. The immunology of asthma. Nat Immunol. 2015;16(1):45–56. Epub 2014/12/19. https://doi.org/10.1038/ni.3049. PubMed PMID: 25521684.

6. Okada H, Kuhn C, Feillet H, Bach JF. The ‘hygiene hypothesis’ for autoimmune and allergic diseases: an update. Clin Exp Immunol. 2010;160(1):1–9. Epub 2010/04/27. https://doi.org/10.1111/j.1365-2249.2010.04139.x. PubMed PMID: 20415844; PubMed Central PMCID: PMCPMC2841828.

7. Herbert O, Barnetson RS, Weninger W, Kramer U, Behrendt H, Ring J. Western lifestyle and increased prevalence of atopic diseases: an example from a small papua new guinean island. World Allergy Organ J. 2009;2(7):130–7. Epub 2009/07/01. https://doi.org/10.1097/WOX.0b013e3181accf27. PubMed PMID: 23283062; PubMed Central PMCID: PMCPMC3650959.

8. Stiemsma LT, Turvey SE. Asthma and the microbiome: defining the critical window in early life. Allergy Asthma Clin Immunol. 2017;13:3. Epub 2017/01/13. https://doi.org/10.1186/s13223-016-0173-6. PubMed PMID: 28077947; PubMed Central PMCID: PMCPMC5217603.

9. Yang JQ, Zhou Y, Singh RR. Effects of Invariant NKT Cells on Parasite Infections and Hygiene Hypothesis. J Immunol Res. 2016;2016:2395645. Epub 2016/08/27. https://doi.org/10.1155/2016/2395645. PubMed PMID: 27563682; PubMed Central PMCID: PMCPMC4987483.

10. Umetsu DT. Early exposure to germs and the Hygiene Hypothesis. Cell Res. 2012;22(8):1210–1. Epub 2012/04/25. https://doi.org/10.1038/cr.2012.65. PubMed PMID: 22525335; PubMed Central PMCID: PMCPMC3411171.

11. Maizels RM, McSorley HJ. Regulation of the host immune system by helminth parasites. J Allergy Clin Immunol. 2016;138(3):666–75. Epub 2016/08/02. https://doi.org/10.1016/j.jaci.2016.07.007. PubMed PMID: 27476889; PubMed Central PMCID: PMCPMC5010150.

12. Sitcharungsi R, Sirivichayakul C. Allergic diseases and helminth infections. Pathog Glob Health. 2013;107(3):110–5. Epub 2013/05/21. https://doi.org/10.1179/2047773213Y.0000000080. PubMed PMID: 23683364; PubMed Central PMCID: PMCPMC4003587.

13. Maizels RM. Parasitic helminth infections and the control of human allergic and autoimmune disorders. Clin Microbiol Infect. 2016;22(6):481–6. Epub 2016/05/14. https://doi.org/10.1016/j.cmi.2016.04.024. PubMed PMID: 27172808.

14. Qiu S, Fan X, Yang Y, Dong P, Zhou W, Xu Y, et al. Schistosoma japonicum infection downregulates house dust mite-induced allergic airway inflammation in mice. PLoS One. 2017;12(6):e0179565. Epub 2017/06/15. https://doi.org/10.1371/journal.pone.0179565. PubMed PMID: 28614408; PubMed Central PMCID: PMCPMC5470717.

15. Osada Y, Shimizu S, Kumagai T, Yamada S, Kanazawa T. Schistosoma mansoni infection reduces severity of collagen-induced arthritis via down-regulation of pro-inflammatory mediators. Int J Parasitol. 2009;39(4):457–64. Epub 2008/10/07. https://doi.org/10.1016/j.ijpara.2008.08.007. PubMed PMID: 18835272.

16. Janssen L, Silva Santos GL, Muller HS, Vieira AR, de Campos TA, de Paulo Martins V. Schistosome-Derived Molecules as Modulating Actors of the Immune System and Promising Candidates to Treat Autoimmune and Inflammatory Diseases. J Immunol Res. 2016;2016:5267485. Epub 2016/09/17. https://doi.org/10.1155/2016/5267485. PubMed PMID: 27635405; PubMed Central PMCID: PMCPMC5011209.

17. Kuprys-Lipinska I, Kuna P. [Changes in the newest recommendations on Asthma Management and Prevention - GINA Report 2014. What should we pay attention to?]. Pneumonol Alergol Pol. 2014;82(5):393–401. Epub 2014/08/19. https://doi.org/10.5603/PiAP.2014.0051. PubMed PMID: 25133806.

18. Falk N. Allergy and Asthma: Asthma Management. FP Essent. 2018;472:25–9. Epub 2018/08/29. PubMed PMID: 30152671.

19. Al-Ahmad M, Arifhodzic N, Nurkic J, Maher A, Rodriguez-Bouza T, Al-Ahmed N, et al. “Real-life” Efficacy and Safety Aspects of 4-Year Omalizumab Treatment for Asthma. Med Princ Pract. 2018;27(3):260–6. Epub 2018/02/08. https://doi.org/10.1159/000487482. PubMed PMID: 29414831; PubMed Central PMCID: PMCPMC6062694.

20. LoVerde PT. Schistosomiasis. Adv Exp Med Biol. 2019;1154:45–70. Epub 2019/07/13. https://doi.org/10.1007/978-3-030-18616-6_3. PubMed PMID: 31297759.

21. Layland LE, Straubinger K, Ritter M, Loffredo-Verde E, Garn H, Sparwasser T, et al. Schistosoma mansoni-mediated suppression of allergic airway inflammation requires patency and Foxp3+ Treg cells. PLoS Negl Trop Dis. 2013;7(8):e2379. Epub 2013/08/24. https://doi.org/10.1371/journal.pntd.0002379. PubMed PMID: 23967364; PubMed Central PMCID: PMCPMC3744427.

22. van der Vlugt LE, Labuda LA, Ozir-Fazalalikhan A, Lievers E, Gloudemans AK, Liu KY, et al. Schistosomes induce regulatory features in human and mouse CD1d(hi) B cells: inhibition of allergic inflammation by IL-10 and regulatory T cells. PLoS One. 2012;7(2):e30883. Epub 2012/02/22. https://doi.org/10.1371/journal.pone.0030883. PubMed PMID: 22347409; PubMed Central PMCID: PMCPMC3275567.

23. Smits HH, Hammad H, van Nimwegen M, Soullie T, Willart MA, Lievers E, et al. Protective effect of Schistosoma mansoni infection on allergic airway inflammation depends on the intensity and chronicity of infection. J Allergy Clin Immunol. 2007;120(4):932–40. Epub 2007/08/11. https://doi.org/10.1016/j.jaci.2007.06.009. PubMed PMID: 17689595.

24. Obieglo K, Schuijs MJ, Ozir-Fazalalikhan A, Otto F, van Wijck Y, Boon L, et al. Isolated Schistosoma mansoni eggs prevent allergic airway inflammation. Parasite Immunol. 2018;40(10):e12579. Epub 2018/08/15. https://doi.org/10.1111/pim.12579. PubMed PMID: 30107039; PubMed Central PMCID: PMCPMC6175163.

25. Mangan NE, van Rooijen N, McKenzie AN, Fallon PG. Helminth-modified pulmonary immune response protects mice from allergen-induced airway hyperresponsiveness. J Immunol. 2006;176(1):138–47. Epub 2005/12/21. https://doi.org/10.4049/jimmunol.176.1.138. PubMed PMID: 16365404.

26. Pacifico LG, Marinho FA, Fonseca CT, Barsante MM, Pinho V, Sales-Junior PA, et al. Schistosoma mansoni antigens modulate experimental allergic asthma in a murine model: a major role for CD4+ CD25+ Foxp3+ T cells independent of interleukin-10. Infect Immun. 2009;77(1):98–107. Epub 2008/10/01. https://doi.org/10.1128/IAI.00783-07. PubMed PMID: 18824533; PubMed Central PMCID: PMCPMC2612239.

27. Medeiros M, Jr., Figueiredo JP, Almeida MC, Matos MA, Araujo MI, Cruz AA, et al. Schistosoma mansoni infection is associated with a reduced course of asthma. J Allergy Clin Immunol. 2003;111(5):947–51. Epub 2003/05/14. https://doi.org/10.1067/mai.2003.1381. PubMed PMID: 12743556.

28. Zhang W, Li L, Zheng Y, Xue F, Yu M, Ma Y, et al. Schistosoma japonicum peptide SJMHE1 suppresses airway inflammation of allergic asthma in mice. J Cell Mol Med. 2019;23(11):7819–29. Epub 2019/09/10. https://doi.org/10.1111/jcmm.14661. PubMed PMID: 31496071; PubMed Central PMCID: PMCPMC6815837.

29. Persson C. In vivo observations provide insight into roles of eosinophils and epithelial cells in asthma. Eur Respir J. 2019;54(4). Epub 2019/06/30. https://doi.org/10.1183/13993003.00470-2019. PubMed PMID: 31248957.

30. Galli SJ, Tsai M. IgE and mast cells in allergic disease. Nat Med. 2012;18(5):693–704. Epub 2012/05/09. https://doi.org/10.1038/nm.2755. PubMed PMID: 22561833; PubMed Central PMCID: PMCPMC3597223.

31. Tsai YG, Niu DM, Yang KD, Hung CH, Yeh YJ, Lee CY, et al. Functional defects of CD46-induced regulatory T cells to suppress airway inflammation in mite allergic asthma. Lab Invest. 2012;92(9):1260–9. Epub 2012/07/04. https://doi.org/10.1038/labinvest.2012.86. PubMed PMID: 22751347.

32. Purroy C, Fairchild RL, Tanaka T, Baldwin WM, 3rd, Manrique J, Madsen JC, et al. Erythropoietin Receptor-Mediated Molecular Crosstalk Promotes T Cell Immunoregulation and Transplant Survival. J Am Soc Nephrol. 2017;28(8):2377–92. Epub 2017/03/18. https://doi.org/10.1681/ASN.2016101100. PubMed PMID: 28302753; PubMed Central PMCID: PMCPMC5533236.

33. Gehrie E, Van der Touw W, Bromberg JS, Ochando JC. Plasmacytoid dendritic cells in tolerance. Methods Mol Biol. 2011;677:127–47. Epub 2010/10/14. https://doi.org/10.1007/978-1-60761-869-0_9. PubMed PMID: 20941607; PubMed Central PMCID: PMCPMC3721973.

34. Kulkarni N, Meitei HT, Sonar SA, Sharma PK, Mujeeb VR, Srivastava S, et al. CCR6 signaling inhibits suppressor function of induced-Treg during gut inflammation. J Autoimmun. 2018;88:121–30. Epub 2017/11/12. https://doi.org/10.1016/j.jaut.2017.10.013. PubMed PMID: 29126851.

35. Tang C, Kamiya T, Liu Y, Kadoki M, Kakuta S, Oshima K, et al. Inhibition of Dectin-1 Signaling Ameliorates Colitis by Inducing Lactobacillus-Mediated Regulatory T Cell Expansion in the Intestine. Cell Host Microbe. 2015;18(2):183–97. Epub 2015/08/14. https://doi.org/10.1016/j.chom.2015.07.003. PubMed PMID: 26269954.

36. Rauch KS, Hils M, Lupar E, Minguet S, Sigvardsson M, Rottenberg ME, et al. Id3 Maintains Foxp3 Expression in Regulatory T Cells by Controlling a Transcriptional Network of E47, Spi-B, and SOCS3. Cell Rep. 2016;17(11):2827–36. Epub 2016/12/16. https://doi.org/10.1016/j.celrep.2016.11.045. PubMed PMID: 27974197.

37. Naval-Macabuhay I, Casanova V, Navarro G, Garcia F, Leon A, Miralles L, et al. Adenosine deaminase regulates Treg expression in autologous T cell-dendritic cell cocultures from patients infected with HIV-1. J Leukoc Biol. 2016;99(2):349–59. Epub 2015/08/28. https://doi.org/10.1189/jlb.3A1214-580RR. PubMed PMID: 26310829.

38. Cheng HY, Gaddis DE, Wu R, McSkimming C, Haynes LD, Taylor AM, et al. Loss of ABCG1 influences regulatory T cell differentiation and atherosclerosis. J Clin Invest. 2016;126(9):3236–46. Epub 2016/08/03. https://doi.org/10.1172/JCI83136. PubMed PMID: 27482882; PubMed Central PMCID: PMCPMC5004951.

39. Zhou Y, Chen H, Liu L, Yu X, Sukhova GK, Yang M, et al. Cathepsin K Deficiency Ameliorates Systemic Lupus Erythematosus-like Manifestations in Fas(lpr) Mice. J Immunol. 2017;198(5):1846–54. Epub 2017/01/18. https://doi.org/10.4049/jimmunol.1501145. PubMed PMID: 28093526; PubMed Central PMCID: PMCPMC5321845.

40. Yan X, Wu C, Chen T, Santos MM, Liu CL, Yang C, et al. Cathepsin S inhibition changes regulatory T-cell activity in regulating bladder cancer and immune cell proliferation and apoptosis. Mol Immunol. 2017;82:66–74. Epub 2016/12/30. https://doi.org/10.1016/j.molimm.2016.12.018. PubMed PMID: 28033540.

41. Liu W, Li H, Zhang X, Wen D, Yu F, Yang S, et al. Prostaglandin I2-IP signalling regulates human Th17 and Treg cell differentiation. Prostaglandins Leukot Essent Fatty Acids. 2013;89(5):335–44. Epub 2013/09/17. https://doi.org/10.1016/j.plefa.2013.08.006. PubMed PMID: 24035274.

42. Jing Y, Kang D, Liu L, Huang H, Chen A, Yang L, et al. Dedicator of cytokinesis protein 2 couples with lymphoid enhancer-binding factor 1 to regulate expression of CD21 and B-cell differentiation. J Allergy Clin Immunol. 2019;144(5):1377–90 e4. Epub 2019/08/14. https://doi.org/10.1016/j.jaci.2019.05.041. PubMed PMID: 31405607.

43. Low MSY, Brodie EJ, Fedele PL, Liao Y, Grigoriadis G, Strasser A, et al. IRF4 Activity Is Required in Established Plasma Cells to Regulate Gene Transcription and Mitochondrial Homeostasis. Cell Rep. 2019;29(9):2634–45 e5. Epub 2019/11/28. https://doi.org/10.1016/j.celrep.2019.10.097. PubMed PMID: 31775034.

44. Croker BA, Tarlinton DM, Cluse LA, Tuxen AJ, Light A, Yang FC, et al. The Rac2 guanosine triphosphatase regulates B lymphocyte antigen receptor responses and chemotaxis and is required for establishment of B-1a and marginal zone B lymphocytes. J Immunol. 2002;168(7):3376–86. Epub 2002/03/22. https://doi.org/10.4049/jimmunol.168.7.3376. PubMed PMID: 11907095.

45. de Oliveira FL, Dos Santos SN, Ricon L, da Costa TP, Pereira JX, Brand C, et al. Lack of galectin-3 modifies differentially Notch ligands in bone marrow and spleen stromal cells interfering with B cell differentiation. Sci Rep. 2018;8(1):3495. Epub 2018/02/24. https://doi.org/10.1038/s41598-018-21409-7. PubMed PMID: 29472568; PubMed Central PMCID: PMCPMC5823902.

46. Gu Y, Jensen PE, Chen X. Immunodeficiency and autoimmunity in H2-O-deficient mice. J Immunol. 2013;190(1):126–37. Epub 2012/12/05. https://doi.org/10.4049/jimmunol.1200993. PubMed PMID: 23209323.

47. Peng C, Eckhardt LA. Role of the Igh intronic enhancer Emu in clonal selection at the pre-B to immature B cell transition. J Immunol. 2013;191(8):4399–411. Epub 2013/09/24. https://doi.org/10.4049/jimmunol.1301858. PubMed PMID: 24058175; PubMed Central PMCID: PMCPMC3810302.

48. Scheikl T, Reis B, Pfeffer K, Holzmann B, Beer S. Reduced notch activity is associated with an impaired marginal zone B cell development and function in Sly1 mutant mice. Mol Immunol. 2009;46(5):969–77. Epub 2008/10/28. https://doi.org/10.1016/j.molimm.2008.09.023. PubMed PMID: 18950867.

49. Flach H, Rosenbaum M, Duchniewicz M, Kim S, Zhang SL, Cahalan MD, et al. Mzb1 protein regulates calcium homeostasis, antibody secretion, and integrin activation in innate-like B cells. Immunity. 2010;33(5):723–35. Epub 2010/11/26. https://doi.org/10.1016/j.immuni.2010.11.013. PubMed PMID: 21093319; PubMed Central PMCID: PMCPMC3125521.

50. Rock JR, Futtner CR, Harfe BD. The transmembrane protein TMEM16A is required for normal development of the murine trachea. Dev Biol. 2008;321(1):141–9. Epub 2008/07/01. https://doi.org/10.1016/j.ydbio.2008.06.009. PubMed PMID: 18585372.

51. Sato T, Okumura F, Ariga T, Hatakeyama S. TRIM6 interacts with Myc and maintains the pluripotency of mouse embryonic stem cells. J Cell Sci. 2012;125(Pt 6):1544–55. Epub 2012/02/14. https://doi.org/10.1242/jcs.095273. PubMed PMID: 22328504.

52. Nuttall RK, Sampieri CL, Pennington CJ, Gill SE, Schultz GA, Edwards DR. Expression analysis of the entire MMP and TIMP gene families during mouse tissue development. FEBS Lett. 2004;563(1-3):129–34. Epub 2004/04/06. https://doi.org/10.1016/S0014-5793(04)00281-9. PubMed PMID: 15063736.

53. Khurana S, George SP. Regulation of cell structure and function by actin-binding proteins: villin’s perspective. FEBS Lett. 2008;582(14):2128–39. Epub 2008/03/01. https://doi.org/10.1016/j.febslet.2008.02.040. PubMed PMID: 18307996; PubMed Central PMCID: PMCPMC2680319.

54. Mehalow AK, Kameya S, Smith RS, Hawes NL, Denegre JM, Young JA, et al. CRB1 is essential for external limiting membrane integrity and photoreceptor morphogenesis in the mammalian retina. Hum Mol Genet. 2003;12(17):2179–89. Epub 2003/08/14. https://doi.org/10.1093/hmg/ddg232. PubMed PMID: 12915475.

55. Harnett MM, Harnett W. Can Parasitic Worms Cure the Modern World’s Ills? Trends Parasitol. 2017;33(9):694–705. Epub 2017/06/14. https://doi.org/10.1016/j.pt.2017.05.007. PubMed PMID: 28606411.

56. Bach JF. The hygiene hypothesis in autoimmunity: the role of pathogens and commensals. Nat Rev Immunol. 2018;18(2):105–20. Epub 2017/10/17. https://doi.org/10.1038/nri.2017.111. PubMed PMID: 29034905.

57. de Ruiter K, Tahapary DL, Sartono E, Soewondo P, Supali T, Smit JWA, et al. Helminths, hygiene hypothesis and type 2 diabetes. Parasite Immunol. 2017;39(5). Epub 2016/12/08. https://doi.org/10.1111/pim.12404. PubMed PMID: 27925245.

58. Maizels RM, McSorley HJ, Smyth DJ. Helminths in the hygiene hypothesis: sooner or later? Clin Exp Immunol. 2014;177(1):38–46. Epub 2014/04/23. https://doi.org/10.1111/cei.12353. PubMed PMID: 24749722; PubMed Central PMCID: PMCPMC4089153.

59. Capron M. Effect of parasite infection on allergic disease. Allergy. 2011;66 Suppl 95:16–8. Epub 2011/06/28. https://doi.org/10.1111/j.1398-9995.2011.02624.x. PubMed PMID: 21668844.

60. Rheinberg CE, Mone H, Caffrey CR, Imbert-Establet D, Jourdane J, Ruppel A. Schistosoma haematobium, S. intercalatum, S. japonicum, S. mansoni, and S. rodhaini in mice: relationship between patterns of lung migration by schistosomula and perfusion recovery of adult worms. Parasitol Res. 1998;84(4):338–42. Epub 1998/05/06. https://doi.org/10.1007/s004360050407. PubMed PMID: 9569102.

61. Cooper PJ. Interactions between helminth parasites and allergy. Curr Opin Allergy Clin Immunol. 2009;9(1):29–37. Epub 2008/12/25. https://doi.org/10.1097/ACI.0b013e32831f44a6. PubMed PMID: 19106698; PubMed Central PMCID: PMCPMC2680069.

62. Baru AM, Hartl A, Lahl K, Krishnaswamy JK, Fehrenbach H, Yildirim AO, et al. Selective depletion of Foxp3+ Treg during sensitization phase aggravates experimental allergic airway inflammation. Eur J Immunol. 2010;40(8):2259–66. Epub 2010/06/15. https://doi.org/10.1002/eji.200939972. PubMed PMID: 20544727.

63. Gabet S, Ranciere F, Just J, de Blic J, Lezmi G, Amat F, et al. Asthma and allergic rhinitis risk depends on house dust mite specific IgE levels in PARIS birth cohort children. World Allergy Organ J. 2019;12(9):100057. Epub 2019/10/24. https://doi.org/10.1016/j.waojou.2019.100057. PubMed PMID: 31641405; PubMed Central PMCID: PMCPMC6796773.

64. Zaccone P, Burton O, Miller N, Jones FM, Dunne DW, Cooke A. Schistosoma mansoni egg antigens induce Treg that participate in diabetes prevention in NOD mice. Eur J Immunol. 2009;39(4):1098–107. Epub 2009/03/18. https://doi.org/10.1002/eji.200838871. PubMed PMID: 19291704.

65. Haeberlein S, Obieglo K, Ozir-Fazalalikhan A, Chaye MAM, Veninga H, van der Vlugt L, et al. Schistosome egg antigens, including the glycoprotein IPSE/alpha-1, trigger the development of regulatory B cells. PLoS Pathog. 2017;13(7):e1006539. Epub 2017/07/29. https://doi.org/10.1371/journal.ppat.1006539. PubMed PMID: 28753651; PubMed Central PMCID: PMCPMC5550006.

66. Liu M, Ju C, Du XF, Shen HM, Wang JP, Li J, et al. Proteomic Analysis on Cercariae and Schistosomula in Reference to Potential Proteases Involved in Host Invasion of Schistosoma japonicum Larvae. J Proteome Res. 2015;14(11):4623–34. Epub 2015/09/16. https://doi.org/10.1021/acs.jproteome.5b00465. PubMed PMID: 26370134.

67. Hansell E, Braschi S, Medzihradszky KF, Sajid M, Debnath M, Ingram J, et al. Proteomic analysis of skin invasion by blood fluke larvae. PLoS Negl Trop Dis. 2008;2(7):e262. Epub 2008/07/17. https://doi.org/10.1371/journal.pntd.0000262. PubMed PMID: 18629379; PubMed Central PMCID: PMCPMC2467291.

68. Li R, Cheng C, Chong SZ, Lim AR, Goh YF, Locht C, et al. Attenuated Bordetella pertussis BPZE1 protects against allergic airway inflammation and contact dermatitis in mouse models. Allergy. 2012;67(10):1250–8. Epub 2012/08/23. https://doi.org/10.1111/j.1398-9995.2012.02884.x. PubMed PMID: 22909095.

69. Chang EE, Yen CM. Eosinophil chemoattracted by eotaxin from cerebrospinal fluid of mice infected with Angiostrongylus cantonensis assayed in a microchamber. Kaohsiung J Med Sci. 2004;20(5):209–15. Epub 2004/07/06. https://doi.org/10.1016/s1607-551x(09)70108-1. PubMed PMID: 15233231.

70. Hopfenspirger MT, Agrawal DK. Airway hyperresponsiveness, late allergic response, and eosinophilia are reversed with mycobacterial antigens in ovalbumin-presensitized mice. J Immunol. 2002;168(5):2516–22. Epub 2002/02/23. https://doi.org/10.4049/jimmunol.168.5.2516. PubMed PMID: 11859146.

71. Setiady YY, Coccia JA, Park PU. In vivo depletion of CD4+FOXP3+ Treg cells by the PC61 anti-CD25 monoclonal antibody is mediated by FcgammaRIII+ phagocytes. Eur J Immunol. 2010;40(3):780–6. Epub 2009/12/30. https://doi.org/10.1002/eji.200939613. PubMed PMID: 20039297.

72. Anders S, Pyl PT, Huber W. HTSeq--a Python framework to work with high-throughput sequencing data. Bioinformatics. 2015;31(2):166–9. Epub 2014/09/28. https://doi.org/10.1093/bioinformatics/btu638. PubMed PMID: 25260700; PubMed Central PMCID: PMCPMC4287950.

73. Anders S, Huber W. Differential expression analysis for sequence count data. Genome Biol. 2010;11(10):R106. Epub 2010/10/29. https://doi.org/10.1186/gb-2010-11-10-r106. PubMed PMID: 20979621; PubMed Central PMCID: PMCPMC3218662.

74. Yu G, Wang LG, Han Y, He QY. clusterProfiler: an R package for comparing biological themes among gene clusters. OMICS. 2012;16(5):284–7. Epub 2012/03/30. Epub 2008/8/19. https://doi.org/10.1089/omi.2011.0118. PubMed PMID: 22455463; PubMed Central PMCID: PMCPMC3339379.

75. Keir ME, Butte MJ, Freeman GJ. & Sharpe AH. PD-1 and its ligands in tolerance and immunity. Annu. Rev. Immunol. 2008; 26, 677–704. Epub 2008/01/05. https://doi.org/10.1146/annurev.immunol.26.021607.090331.

76. Yu A, Zhu L, Altman NH. & Malek TR. A low interleukin-2 receptor signaling threshold supports the development and homeostasis of T regulatory cells. Immunity. 2009; 30, 204–17. Epub 2009/02/03. https://doi.org/10.1016/j.immuni.2008.11.014. PMCID: PMC2962446.

77. Henderson JG. & Hawiger D. Regulation of extrathymic Treg cell conversion by CD5. Oncotarget. 2015; 6, 26554–26555. Epub 2015/10/10. https://doi.org/10.18632/oncotarget.5809. PMCID: PMC4694933.

78. Watanabe T, Masuyama J, Sohma Y, Inazawa H, Horie K, Kojima K, et al. CD52 is a novel costimulatory molecule for induction of CD4+ regulatory T cells. Clin. Immunol. 2006; 120, 247–259. Epub 2006/06/27. https://doi.org/10.1016/j.clim.2006.05.006.

79. Massoud AH, Yona M, Xue D, Chouiali F, Alturaihi H, Ablona A, Mourad W, Piccirillo CA, Mazer BD. Dendritic cell immunoreceptor: a novel receptor for intravenous immunoglobulin mediates induction of regulatory T cells. J. Allergy Clin. Immunol. 2014; 133, 853–863 e5. Epub 2013/11/12. https://doi.org/10.1016/j.jaci.2013.09.029.

80. Kudo-Saito C, Shirako H, Ohike M, Tsukamoto N & Kawakami Y. CCL2 is critical for immunosuppression to promote cancer metastasis. Clin. Exp. Metastasis. 2013; 30, 393–405. Epub 2012/11/13. https://doi.org/10.1007/s10585-012-9545-6.

